# Systemic MIF facilitates chronic lymphocytic leukemia development independent of its cellular source

**DOI:** 10.1101/2025.02.22.639632

**Authors:** Viktoria Kohlhas, Nina Reinart, Natascha Rosen, Sebastian Reinartz, Phuong-Hien Nguyen, Michael Hallek

**Affiliations:** University of Cologne, Faculty of Medicine and University Hospital Cologne, Department I of Internal Medicine, Center for Integrated Oncology Aachen Bonn Cologne Duesseldorf; Center for Molecular Medicine Cologne; CECAD Center of Excellence on Cellular Stress Responses in Aging-Associated Diseases, 50931 Cologne, Germany

**Keywords:** CLL, MIF, conditional knockout, TCL1 transgenic mice, B cells, macrophages

## Abstract

Macrophage migration inhibitory factor (MIF) is broadly produced by various cell types, particularly immune cells, and functions as a key modulator of innate and adaptive immunity. Increasing evidence has linked MIF to the pathogenesis of both solid tumors and hematologic malignancies, including chronic lymphocytic leukemia (CLL). We previously showed that the global deletion of Mif in the *TCL1* transgenic mouse model for CLL significantly delayed disease development leading to longer overall survival of the knockout mice. In this study, we demonstrated that adaptive transfer of murine CLL cells failed to establish disease in Mif-deficient recipients due to impaired homing of leukemic cells into the spleens, indicating that host-derived Mif is essential for leukemic infiltration and expansion. To identify the most relevant source of Mif in CLL, we generated two CLL mouse strains with B-lymphoid- or myeloid-lineage-specific Mif deletion. In contrast to the global Mif knockout, neither conditional Mif knockout significantly altered CLL progression, illustrating that the cellular source of Mif is less critical than its systemic presence in the tissue environment. Taken together, these in vivo findings indicate that MIF plays a relevant role in CLL pathogenesis, acting independently of its specific cellular origin.

**Highlights:**

- CLL cells failed to establish disease in Mif-deficient hosts due to impaired tissue homing
- Conditional deletion of Mif in B lymphoid- or myeloid cells did not significantly impact CLL progression in vivo
- Systemic MIF is critical for CLL pathogenesis, independent of its cellular source

## Introduction

Since the discovery of macrophage migration inhibitory factor (MIF) as the first cytokine to inhibit the migration of macrophages (Bloom and Bennett 1966; David 1966), MIF has been shown to be a pleiotropic modulator of innate and adaptive immune responses. MIF is produced and released by almost all cell types, including macrophages (T. Calandra et al. 1994), monocytes (Flieger et al. 2003), T and B lymphocytes (Bacher et al. 1996), eosinophils (Rossi et al. 1998), neutrophils (Daryadel et al. 2006) and mast cells (H. Chen et al. 1998). Binding of MIF to its receptor, the CD74-CD44 complex, activates the MAPK-ERK and PI3K-AKT pathways and induces NFκB activation and production of pro-inflammatory target genes, promoting cell survival and inhibiting apoptosis (Leng et al. 2003; Shi et al. 2006); (Thierry Calandra and Roger 2003). MIF also binds to chemokine receptors, including CXCR2, CXCR4, and CXCR7, thereby stimulating cell migration, chemotaxis, and immune responses (Bernhagen et al. 2007; Weber et al. 2008; Alampour-Rajabi et al. 2015). MIF activates monocytes and macrophages and optimizes their expression of TNF-α, IL-1, and PGE2 (Brennan-Bourdon et al. 2015; Cao et al. 2005; Yaddanapudi et al. 2016). In B cells, activation of the surface receptor complex CD74/CD44 by MIF induces the production of IL-8 and increases the resistance to apoptosis via upregulation of BCL-2 (Binsky et al. 2007; Gore et al. 2008). MIF may also influence other pathways, such as TP53 (Fingerle-Rowson et al. 2003; Jung, Seong, and Ha 2008) and JAB1-AP1 (Kleemann et al. 2000).

MIF plays a significant role as a proinflammatory and immuno-regulatory cytokine during inflammatory responses, both under physiological and pathological conditions such as infectious- and autoimmune diseases (Calandra et al. 1994; Kang and Bucala 2019; Sumaiya et al. 2022). Mif-knockout mice develop without apparent deficits under laboratory conditions (Fingerle-Rowson et al. 2003). Upon pathogen challenge, Mif-knockout mice exhibit alleviated inflammatory responses and improved outcomes due to protection against cytokine storms and organ damage (Bozza et al. 1999; Smith et al. 2019; Jinhong Li et al. 2018; Koebernick et al. 2002), underscoring its role in modulating the magnitude of the immune response.

Given the role of MIF in inflammation and putative signaling related to tumorigenesis, there is a growing recognition that MIF is involved in the pathogenesis of various cancers (Bucala and Donnelly 2007); (Valdez et al. 2024), including solid tumors like non-small cell lung cancer (Howard et al. 2004), prostatic adenocarcinoma (Meyer-Siegler and Hudson 1996), bladder cancer (Taylor et al. 2007), hepatocellular carcinoma (Liao et al. 2023) and melanoma (Tanese and Ogata 2024); as well as in several hematologic malignancies including myeloproliferative neoplasm (Pritchard et al. 2024), acute myeloid leukemia (Spertini et al. 2024) B cell lymphoma (Talos et al. 2005), and chronic lymphocytic leukemia (CLL) (Reinart et al. 2013). In most malignant conditions, MIF has been characterized as a tumor-promoting factor that regulates both intrinsic oncogenic signaling pathways within tumor cells and the composition and dynamics of the tumor immune microenvironment (Valdez et al. 2024).

CLL is the most frequent leukemia in the Western Hemisphere. It is a monoclonal B-cell disorder characterized by the expansion and accumulation of morphologically mature but functionally impaired lymphocytes (Hallek et al. 2018). The apoptosis resistance and survival of CLL cells depend highly on interactions with the tumor microenvironment (TME), rendering it a disease “addicted to the host” (Jan A. Burger and Gribben 2014). In CLL, the TME comprises cellular components such as macrophages, fibroblasts, T cells, and soluble molecules like chemokines and cytokines (Vom Stein, Hallek, and Nguyen 2023; Jestrabek et al. 2024).

When intercrossed with the *TCL1* mouse model for CLL (Bichi et al. 2002; Koch et al. 2020), Mif-knockout CLL mice showed a delayed onset of leukemia, reduced splenomegaly and hepatomegaly, and a significantly prolonged survival than *TCL1*^*tg/wt*^ controls (Reinart et al. 2013). Mif-deficient CLL cells were more susceptible to apoptosis *in vitro*. By using neutralizing anti-Mif antibodies, the survival of CLL cells was drastically reduced on a macrophage feeder layer. The number of macrophages was significantly reduced in the spleen and bone marrow of *TCL1*^*tg/wt*^ *Mif*^*-/-*^ mice. Additionally, the migratory activity of *TCL1*^*tg/wt*^*Mif*^*-/-*^ macrophages decreased in comparison to *TCL1*^*tg/wt*^ *Mif*^*wt/wt*^ macrophages. These results indicated that MIF supported the development of CLL *in vivo* by enhancing the interaction of CLL cells with macrophages. In CLL patient samples, MIF mRNA and protein levels were significantly increased in CLL cells compared to B cells of healthy donors. Additionally, MIF plasma levels were strongly elevated (Reinart et al. 2013). The MIF receptors CXCR4 and CD74 are upregulated on CLL cells and are pivotal for leukemic cell viability and migration, cooperatively promoting the survival and homing of CLL cells upon MIF stimulation (J. A. Burger, Burger, and Kipps 1999; Binsky et al. 2006; Thavayogarajah et al. 2022). Although CD74 appears to be dispensable for CLL progression in the *TCL1* mouse model (Barthel et al. 2020), the absence of its coreceptor CD44 results in a longer survival of CLL-diseased mice via MCL1 modulation (Fedorchenko et al. 2013), suggesting that Mif exerts its leukemia-promoting effects at least in part through this complex *in vivo*.

As MIF is a broadly expressed cytokine, it remains unclear which cell type is relevant for the secretion and the disease-promoting effects of MIF in CLL. To address this question, we generated two mouse strains with B lymphoid- and myeloid-lineage-specific deletion of Mif to identify the relevant source of MIF in CLL.

## Materials and Methods

### Mice

All mouse studies were approved by the state authorities of North Rhine-Westphalia, Germany (Landesamt für Natur-, Umwelt-und Verbraucherschutz Nordrhein-Westfalen (LANUV)), approvals #84-02.04.2014.A146 #81-02.04.2019.A009 and #84-02.04.2016.A058. *Eµ-TCL1* transgenic mice (Bichi et al. 2002) were crossed with *Mif*^*fl/fx*^ mice (Brocks et al. 2017). For conditional Mif knockout in B cells and myeloid lineage, *Eµ-TCL1*^*tg/wt*^ *Mif*^*fl/fl*^ mice were crossed with either *CD19*^*cre/wt*^ (Rickert, Roes, and Rajewsky 1997) or *LysM*^*cre/wt*^ (Clausen et al. 1999) mice, respectively. Husbandry, procedures for blood and tissue sample collection, determination of overall survival, and differential blood counts were performed as described previously (Nguyen et al. 2016; Kohlhas, Hallek, and Nguyen 2020).

### RNA-isolation and quantitative RT-PCR

RNA was isolated using Trizol reagent (Invitrogen). Reverse transcription was performed with the ProtoScriptII-First Strand cDNA Synthesis Kit (New England Biolabs). Quantitative real-time PCR for MIF was run on a Light Cycler 480 system (Roche Diagnostics) using TaqMan Fast Universal PCR Master Mix (Applied Biosystems (ThermoFisher Scientific). mRNA levels were normalized to peptidylprolyl isomerase A (PPIA).

Primer: Mm02342430_g1 Ppia FAM PN4453320, Mm01611157_gH Mif, Gm163+ FAM PN4453320

### Fluorescence-activated cell sorting (FACS)

Splenocytes of five wild type (C57BL/6) mice were sorted by FACS into T cells (CD3-pacific blue), NK cells (NK1.1-Alexa Fluor 488), B cells (CD19-APC-Cy7), macrophages (F4/80-PE), and endothelial cells (CD146-APC).

### Syngeneic transplantation of murine CLL cells

Transplantation of murine CLL cells was previously described (Nguyen et al. 2016; Koch et al. 2020). All donor- and recipient mice were backcrossed for over 10 generations to a C57BL/6J genetic background. Freshly homogenized and filtered splenocytes from moribund *Eµ-TCL1* mice were layered with the Lymphocyte Separate Medium (PAA), followed by centrifugation and separation of interphase-concentrated mononuclear cells. After several washing steps, CLL cells were viable, frozen at -150 °C, and thawed upon transplantation. *Eµ-TCL1* CLL cells were thawed and injected intraperitoneally into age- and sex-matched WT (n = 6) and *Mif*^*-/-*^ (n = 9) mice. Blood was drawn every 2 weeks and analyzed for CD5^+^ CD19^+^ positive cells to monitor CLL development.

### *In vivo* short-term homing assay

Isolated *Eµ-TCL1* CLL cells (n = 2) were stained with CFSE (5 µM, Abcam) and injected into sex-matched WT (n = 4) and Mif^-/-^ (n = 4) recipients intravenously. After 3 hours, mice were sacrificed, and splenocytes and bone marrow cells were isolated. Fc-Receptor was blocked with FcR Blocking Reagent (Miltenyi Biotec), and leukocytes were stained with an anti-CD45-VioBlue (Miltenyi Biotec) antibody. For intracellular staining of TCL1 (TCL1(human)-Alexa Fluor 647), the IntraPrep Permeabilization Reagent Kit (Beckman Coulter) was used according to the manufacturer’s instructions. Labeled samples were run on a MACSQuant X (Miltenyi Biotec) flow cytometer, and data were analyzed using Kaluza 2.0 Flow Analysis Software (Beckman Coulter).

## Results

Global Mif deletion in the *TCL1*^*tg/wt*^ mice resulted in a significantly delayed development of CLL due to defective homing of CLL cells to the lymphoid organs essential for leukemic cell expansion (Reinart et al. 2013). To determine whether the absence of Mif in the tumor microenvironment impaired CLL development, *TCL1*^*tg/wt*^ CLL cells were transplanted into wild type (WT) or *Mif*^*-/-*^ mice, and CLL development was monitored in recipients over 21 weeks. While WT animals showed an increasing number of CLL cells in peripheral blood, almost no CLL cells could be detected in the peripheral blood of *Mif*^*-/-*^ animals throughout the monitoring time (Fig. 1A). The median survival of transplanted WT mice was 139 days, whereas none of the transplanted *Mif*^*-/-*^ recipients succumbed to leukemia and did not reach the defined endpoint for this experiment (Fig. 1B). Because homing of transplanted CLL cells to lymphoid organs such as the spleen is crucial for outgrowth of leukemia cells in recipient mice, we tested if CLL cells could migrate to the spleens in short-term homing assays. *TCL1* CLL cells were transplanted into WT and *Mif*^*-/-*^ recipients, and after 3 h, the homing of CLL cells into the spleen was measured via flow cytometry of the *TCL1* transgene. Interestingly, the number of CLL cells was reduced in *Mif*^*-/-*^ recipients (Fig. 1C). These results confirmed our hypothesis that a Mif deficient microenvironment failed to recruit CLL cells into the lymphoid homing organs, thereby preventing CLL growth.

**Figure 1.**
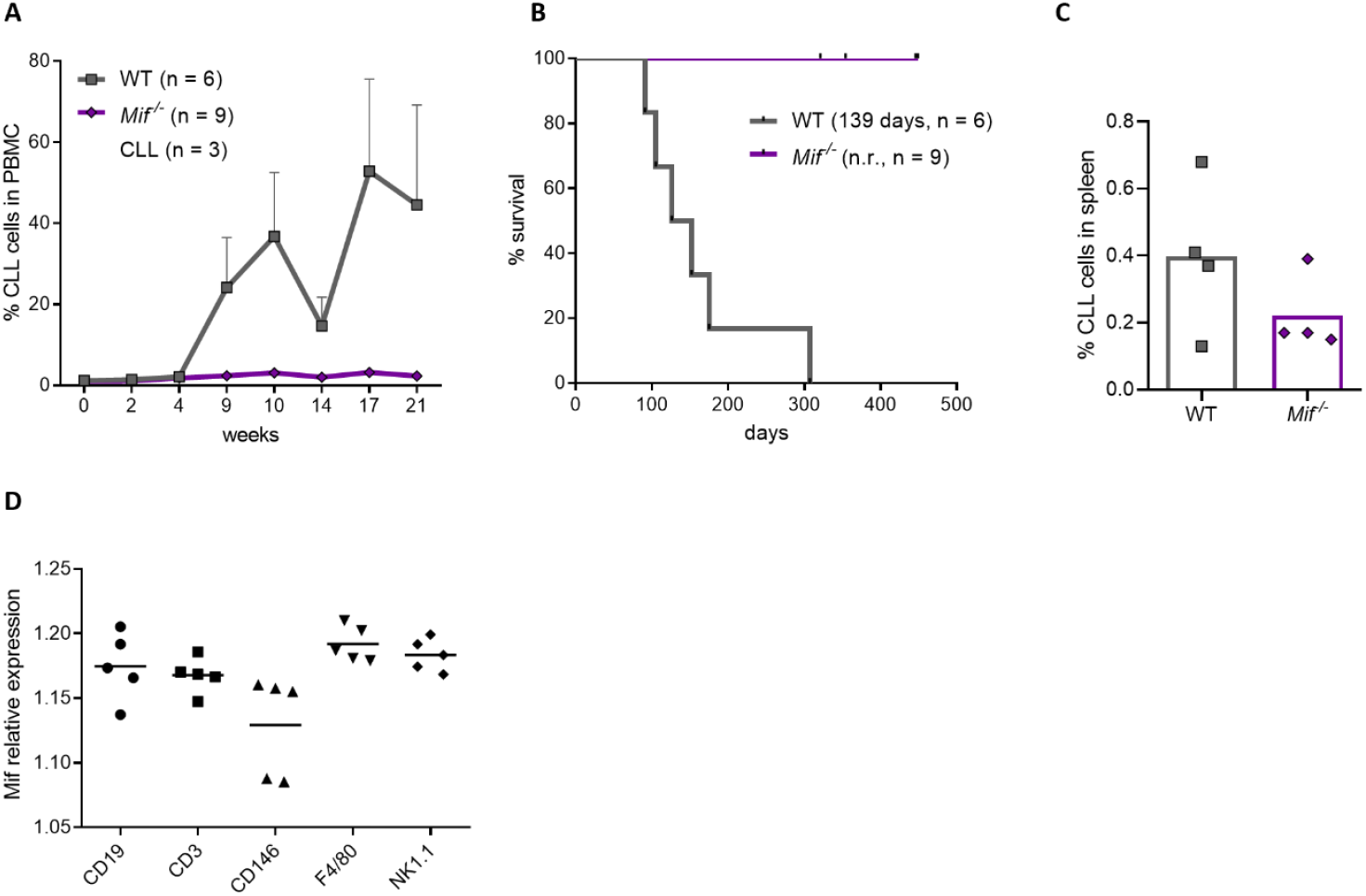
Mif is a critical cytokine for leukemia engraftment in the CLL adoptive transfer model. A: Flow cytometric analysis of CD19^+^ CD5^+^ CLL cells in the peripheral blood of WT and *Mif*^*-/-*^ recipients after syngeneic transplantation of *TCL1*^*tg/wt*^ CLL cells. B: Survival of CLL-transplanted WT and *Mif*^*-/-*^ mice. *Mif*^*-/-*^ recipients did not reach the primary endpoint. C: Homing of CLL cells in spleens of WT and *Mif*^*-/-*^ 3 h after transplantation of *TCL1*^*tg/wt*^ CLL cells. D: Relative Mif RNA expression in isolated cell types of wild type mice. FACS-sorted B cells (CD19), T cells (CD3), endothelial cells (CD146), macrophages (F4/80), and NK cells (NK1.1) from murine spleens were subjected to qRT-PCR.

To uncover the cellular source of Mif in the CLL microenvironment, gene expression of Mif was determined in different purified cell types from the spleens of WT C57BL/6 mice. B cells (CD19^+^), T cells (CD3^+^), endothelial cells (CD146^+^), macrophages (F4/80^+^), and NK cells (NK1.1^+^) were harvested and sorted from murine spleens based on their surface marker expression by FACS. Mif was found to be expressed in all cell types using qRT-PCR. The highest expression was found in macrophages (relative expression: 1.192), followed by NK cells (1.183) and B cells (1.175). A lower expression was detected in T cells (1.168) and the lowest in endothelial cells (1.129) (Fig. 1D).

As Mif was shown to be elevated in CLL cells and highly expressed in the B cell compartment, we sought to determine if Mif expression in the leukemic B cells would be critical for disease development. For this purpose, we generated B cell-specific Mif knockout mice by intercrossing *Mif*^*fl/fl*^ mice with *TCL1*^*tg/wt*^ and *CD19*^*cre/wt*^ mice to obtain a B cell-specific Mif knockout (*TCL1*^*tg/wt*^ *CD19*^*cre/wt*^ *Mif*^*fl/fl*^) (Fig. 2A) and monitored CLL development in blood samples of *TCL1*^*tg/wt*^ *CD19*^*wt/wt*^ *Mif*^*fl/fl*^ (referred to as *TCL1*^*tg/wt*^) and *TCL1*^*tg/wt*^ *CD19*^*cre/wt*^ *Mif*^*fl/fl*^ mice over 12 months. In contrast to the significantly delayed CLL progression observed in global *Mif*^*-/-*^ mice, we observed only moderate reductions of the leukocyte counts of *TCL1*^*tg/wt*^ *CD19*^*cre/wt*^ *Mif*^*fl/fl*^ at month 9 and month 12 (Fig. 2B). In line with these findings, CLL burden determined by the percentage of CD5^+^ CD19^+^ cells in PBMC (Fig. 2C) and CLL count per µl blood (Fig. 2D) showed no significant differences in the B cell-specific Mif knockout compared to controls. There was only a non-significant trend of a lower CLL burden at month 6, suggesting a slight delay in disease progression at the early stage. Additionally, no differences in the weight of spleens (1.57 gr vs. 1.81 gr) and livers (3.54 gr vs.2.97 gr) were observed between both genotypes (Fig. 2E). The B cell-specific Mif knockout did not affect the overall survival of mice without CLL burden, as WT mice lived 827 days (n = 10) and *CD19*^*cre/wt*^ *Mif*^*fl/fl*^ lived 878 days (n = 19) on average. Mice with the *TCL1* transgene lived significantly shorter, as expected. However, no significant difference in the overall survival between *TCL1*^*tg/wt*^ (430 days, n = 10) and *TCL1*^*tg/wt*^ *CD19*^*cre/wt*^ *Mif*^*fl/fl*^ (402 days, n = 22) was observed (Fig. 2F).

**Figure 2.**
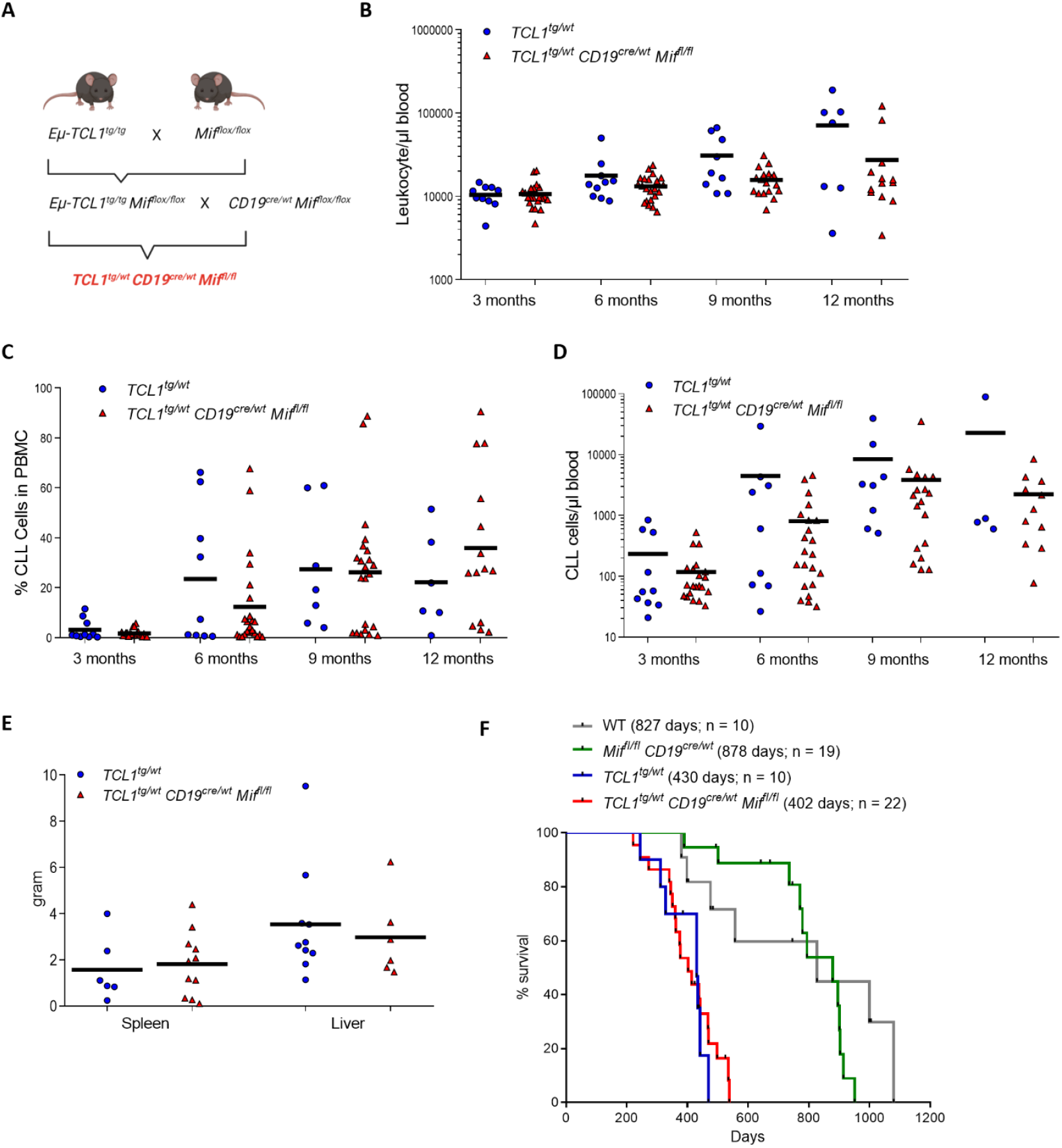
*TCL1*^*tg/wt*^ *CD19*^*cre/wt*^ *Mif*^*fl/fl*^ mice show no clear difference in CLL burden compared with *TCL1*^*tg/wt*^ mice. A: Schematic overview of the breeding strategy to establish the *TCL1*^*tg/wt*^ *CD19*^*cre/wt*^ *Mif*^*fl/fl*^ mouse strain. B: Count of leukocytes in 1 µl blood of *TCL1*^*tg/wt*^ and *TCL1*^*tg/wt*^ *CD19*^*cre/wt*^ *Mif*^*fl/fl*^ over one year. C: Flow cytometric analysis of CD19^+^ CD5^+^ CLL cells in the peripheral blood of *TCL1*^*tg/wt*^ and *TCL1*^*tg/wt*^ *CD19*^*cre/wt*^ *Mif*^*fl/fl*^ mice over one year. D: Count of CLL cells in 1 µl blood of *TCL1*^*tg/wt*^ and *TCL1*^*tg/wt*^ *CD19*^*cre/wt*^ *Mif*^*fl/fl*^ over one year. E: Spleen and liver weights of age-matched and moribund TCL1^tg/wt^ and *TCL1*^*tg/wt*^ *CD19*^*cre/wt*^ *Mif*^*fl/fl*^ mice. F: Kaplan-Meier curves representing the overall survival of WT, *Mif*^*fl/fl*^, *TCL1*^*tg/w*t^, *TCL1*^*tg/wt*^ *CD19*^*cre/wt*^ *Mif*^*fl/fl*^ mice from birth to moribund.

The lacking effect of Mif depletion in B-CLL cells prompted us to examine the impact of Mif depletion in other cell types of the microenvironment. As Mif showed the highest expression in macrophages, and we had observed that a reduction of macrophages in *TCL1*^*tg/wt*^ *Mif*^*-/-*^ hindered CLL expansion (Reinart et al. 2013), we crossed the *TCL1*^*tg/wt*^ *Mif*^*fl/fl*^ mice with *LysM*^*cre/wt*^ mice, which express the Cre recombinase under the control of the myeloid cell-specific lysozyme 2 gene to generate a myeloid-derived cell-specific conditional knockout (Clausen et al. 1999) (Fig. 3A). Again, we monitored CLL burden in *TCL1*^*tg/wt*^ *LysM*^*cre/wt*^ *Mif*^*fl/fl*^ and control mice over 12 months of age. We did not detect any differences in leukocyte count (Fig. 3B), as well as no differences in the parameters reflecting the CLL development in the peripheral blood in the conditional macrophage knockout of Mif (Fig. 3C). Additionally, we did not detect significant differences in the weight of spleens (1.867 g versus 0.785 g) and livers (2.73 g versus 2.111 g) between *TCL1*^*tg/wt*^ and *TCL1*^*tg/wt*^ *LysM*^*cre/wt*^ *Mif*^*fl/fl*^ (Fig. 3D). The overall survival of CLL mice was not affected by the conditional knockout, as *TCL1*^*tg/wt*^ mice lived 436 days (n = 27) and *TCL1*^*tg/wt*^ *LysM*^*cre/wt*^ *Mif*^*fl/fl*^ mice lived 397 days (n = 26) on average (Fig. 3E). Finally, we also did not observe any differences in the percentage of myeloid cells or monocytes in the peripheral blood of *LysM*^*cre/wt*^ and *TCL1*^*tg/wt*^ *LysM*^*cre/wt*^ *Mif*^*fl/fl*^ and their respective controls, WT or *TCL1*^*tg/wt*^ (Supplemental Fig. 1A & B). Overall, the macrophage-specific deletion of Mif did not influence the development or progression of CLL in the *TCL1*^*tg/wt*^ *LysM*^*cre/wt*^ *Mif*^*fl/fl*^ mouse model.

**Fig. 3.**
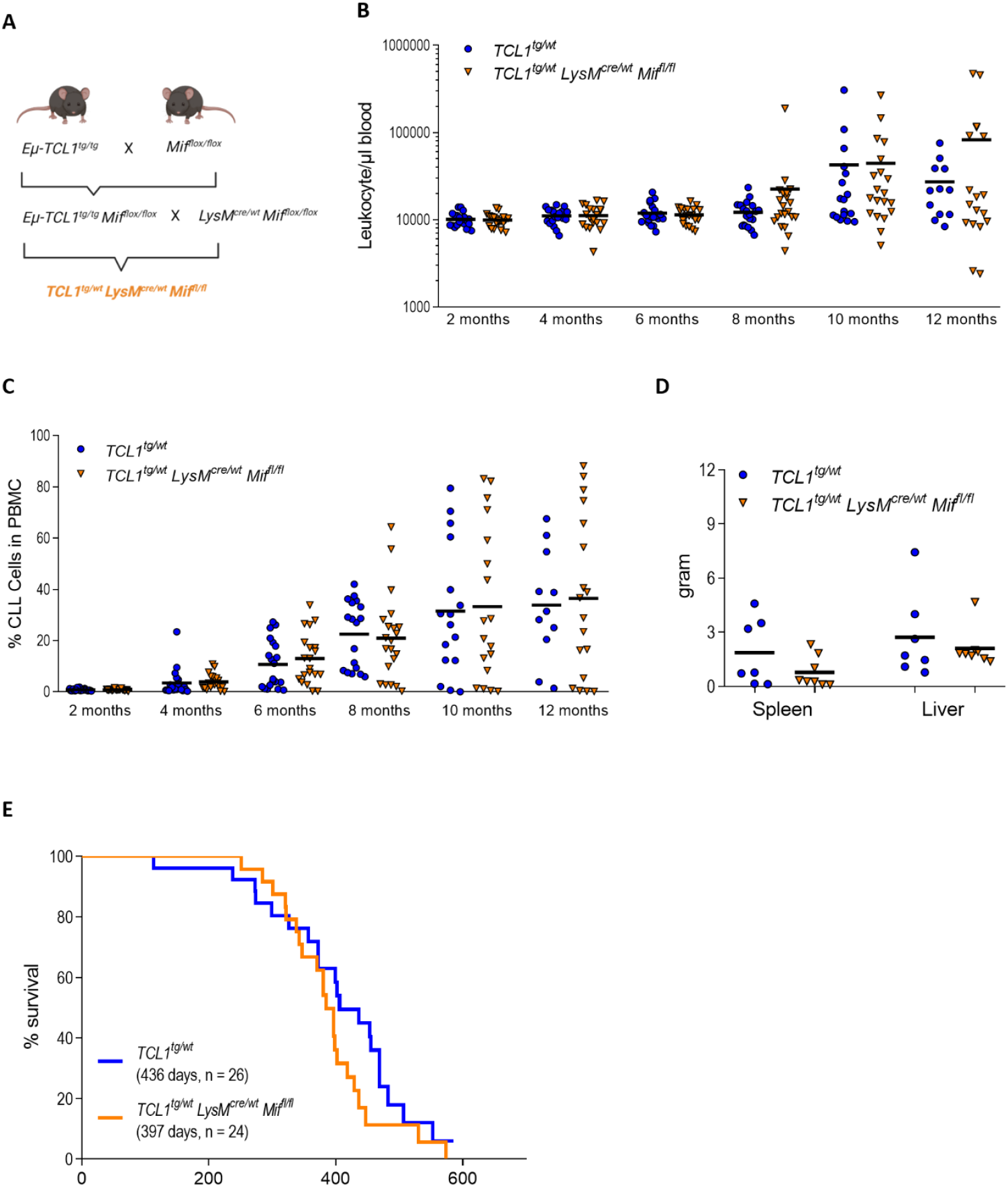
*TCL1*^*tg/wt*^ *LysM*^*cre/wt*^ *Mif*^*fl/fl*^ mice show no difference in CLL burden compared with *TCL1*^*tg/wt*^ mice. A: Schematic overview of the breeding strategy to establish the *TCL1*^*tg/wt*^ *LysM*^*cre/wt*^ *Mif*^*fl/fl*^ mouse strain. B: Count of Leukocytes in 1 µl blood of *TCL1*^*tg/wt*^ and *TCL1*^*tg/wt*^ *LysM*^*cre/wt*^ *Mif*^*fl/fl*^ over one year. C: Flow cytometric analysis of CD19^+^ CD5^+^ CLL cells in the peripheral blood of *TCL1*^*tg/wt*^ and *TCL1*^*tg/wt*^ *LysM*^*cre/wt*^ *Mif*^*fl/fl*^ mice over one year. D: Spleen and liver weights of age-matched, moribund *TCL1*^*tg/wt*^ and *TCL1*^*tg/wt*^ *LysM*^*cre/wt*^ *Mif*^*fl/fl*^ mice. E: Kaplan-Meier curves representing the overall survival of *TCL1*^*tg/wt*^ and *TCL1*^*tg wt*^ *LysM*^*cre/wt*^ *Mif*^*fl/fl*^ mice from birth to moribund.

## Discussion

We previously showed that the absence of the cytokine Mif delayed leukemia development in the *TCL1* mouse model for CLL (Reinart et al. 2013). Mif knockout reduced the survival of CLL cells and the number of macrophages in all typical homing organs of CLL cells. In this study, we demonstrated that CLL cells from *TCL1* mice failed to home to the spleens of *Mif*^*-/-*^ recipient mice, resulting in a lack of CLL outgrowth in these knockout animals. These findings formally confirm that the presence of Mif in the host is highly relevant to recruit CLL cells to the homing organs – a prerequisite for leukemic cell survival and growth. In contrast, there was no significant difference in CLL progression in mice deficient of Mif in B cells or the myeloid lineage. The phenotypes of these two conditional Mif knockout mouse models illustrate that the cellular source of Mif is less critical than the systemic presence of Mif in the tissue environment for CLL development.

CLL cells are highly dependent on various stimuli from their tumor microenvironment, and neither a single cell type nor a soluble factor alone can fully confer malignant potential to CLL cells. Notwithstanding, several cytokines, including CXCL13, BAFF, APRIL, and MIF, have been shown to play crucial roles in CLL initiation and progression (Ullah et al. 2024; J. A. Burger, Burger, and Kipps 1999; Vom Stein, Hallek, and Nguyen 2023). Our *in vivo* transplantation experiment and homing assay underscore the critical role of MIF in recruiting leukemic cells to their homing organs, such as the spleen. Given that the splenic environment in the *TCL1* mouse model mimics the lymph nodes in CLL patients (Collard et al. 2022) – the primary sites for CLL cell proliferation (Mittal et al. 2014) – these findings indicate an essential role of MIF in leukemic cell homing and disease development.

MIF is a ubiquitous cytokine stored in intracellular pools and released by various cell types. Our analysis validates its presence across all cell types in mice, with higher expression in immune cells compared to endothelial cells. The highest expression of Mif in macrophages reflects its crucial function in this lineage, regulating macrophage activation, recruitment, and cytokine production. MIF enhances pro-inflammatory responses while countering the anti-inflammatory effects of glucocorticoids (T. Calandra et al. 1995). Macrophages lacking MIF exhibit reduced activation and migration, dampening immune responses — a beneficial effect in chronic inflammatory diseases (Thierry Calandra and Roger 2003; Thiele, Donnelly, and Mitchell 2022). In cancer, MIF drives macrophage polarization into immunosuppressive, pro-tumorigenic M2-like tumor-associated macrophages (TAMs), which secrete IL-10 and TGF-β while promoting angiogenesis, tumor cell migration, and immune suppression. MIF also facilitates macrophage recruitment to the vicinity of tumors, supporting disease progression. In CLL, global Mif knockout reduces macrophage recruitment to the spleen and bone marrow (Reinart et al. 2013), yet macrophage-specific Mif knockout does not affect disease progression. This observation corroborates that CLL establishment requires the mere presence of Mif, independently of its cellular source. Although we have not investigated the potential influence of Mif depletion in other cell types, such as T cells or NK cells, it seems unlikely that deleting the *Mif* gene in further individual cell types can recapitulate the strong effects observed in the global Mif knockout background.

In healthy B cells, MIF promotes survival, activation, and proliferation during immune processes. However, MIF production becomes dysregulated in CLL, and leukemia cells rely on autocrine MIF signaling, activating NF-κB and BCL2 survival pathways via the CD74 receptor (Binsky et al. 2007; Gore et al. 2008). This intrinsic role of MIF may partially account for the delayed disease progression observed at earlier stages in the B cell specific Mif knockout mice, albeit not significant. Notwithstanding, the lack of Mif production in leukemic B cells alone was insufficient to hinder CLL development, illustrating that soluble Mif released by non-leukemic cells can also efficiently activate the Mif signaling cascade. Unlike solid tumors, where MIF drives local tumor proliferation and immune dysfunction (Wirtz et al. 2021), CLL’s dispersed nature diminishes the impact of autocrine signaling, making systemic MIF more critical. This systemic influence contrasts with hepatocellular carcinoma, where targeting MIF in hepatocytes reduces tumor growth (Wirtz et al. 2021). Given the high expression of MIF and its receptors in CLL cells, it is likely that the low tissue density and the rapid dispersion of cytokines in the blood and lymphoid organs significantly diminish dependence on autocrine signaling. This could explain why targeting Mif production in individual cell types is less effective in CLL than solid tumors, emphasizing the importance of systemic MIF in the leukemic environment.

Given the increasing evidence of MIF’s crucial role in cancer development, MIF has emerged as a promising therapeutic target in various cancers, including CLL. Therapeutic strategies targeting MIF include anti-MIF neutralizing antibodies, MIF antagonists, and small molecule inhibitors. However, MIF presents a challenging target for cancer therapy due to its widespread presence in healthy tissues and circulation. An oxidized form (oxMIF) is linked explicitly with sites of inflammation and solid tumors. Targeting oxMIF with a monoclonal anti-oxMIF antibody showed promising antitumor activity *in vitro* and *in vivo* (Thiele, Donnelly, and Mitchell 2022; Rossmueller et al. 2023). Recently, novel therapeutic approaches have been developed to inhibit MIF activity selectively. Small-molecule allele-selective transcriptional inhibitor of the MIF immune susceptibility locus was generated, increasing the potential for precision-based MIF inhibitors (Jia Li et al. 2024). Furthermore, MIF-directed small molecules have been shown to enhance ferroptosis by impairing DNA repair mechanisms, suggesting a new druggable route for targeting MIF (D. Chen et al. 2024). The first development of potent MIF-directed PROTACs (proteolysis-targeting chimeras) has been reported to induce MIF degradation and disrupt its protein-protein interaction network, leading to anti-proliferative activity in lung cancer cells (Xiao et al. 2021). In CLL, targeting the MIF receptor CD74 with milatuzumab has shown potential in clinical trials (Haran et al. 2018). However, a significant challenge in the treatment of CLL remains the limited options for refractory patients. Novel combination therapies targeting the tumor microenvironment are currently considered promising for these cases (Lewis, Vom Stein, and Hallek 2024). Expanding the range of microenvironmental targets could increase potential treatment options for those patients.

Taken together, our study shows that MIF plays a relevant role in the development of CLL when present in the microenvironment, regardless of the source of production. In the early stages of disease, the presence of Mif in lymphoid tissues is indispensable for the homing of leukemic cells into the lymphoid organs, where CLL cell survival and proliferation mainly occur. CLL cells can effectively employ Mif signaling in both autocrine and non-autocrine manners. This opens potential avenues for using MIF targeting strategies to treat this leukemia.

## Supporting information

Kohlhas et al_Mif in CLL_Supplement

## Author Contribution

V.K. analyzed data and wrote the manuscript. N.R. designed and conducted experiments. Na.R., S.R. conducted experiments. P.-H.N. designed experiments, analyzed data, and wrote the manuscript. M.H. initiated, designed, and supervised the study. All authors read, revised, and approved the manuscript.

