## Supplementary material for "Systemic MIF facilitates chronic lymphocytic leukemia development independent of its cellular source": Kohlhas et al_Mif in CLL_Supplement

**Supplemental Figure 1 and Legend**

**A**

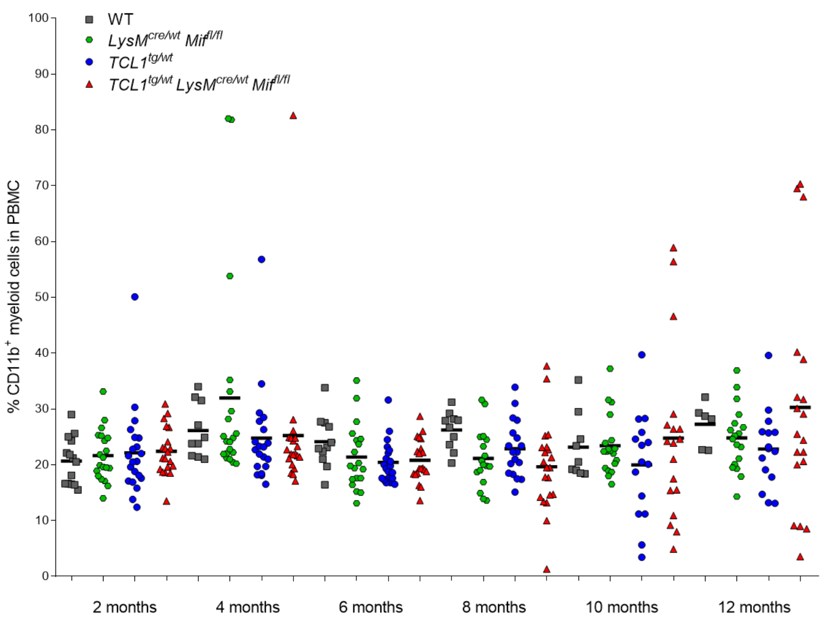


**B**

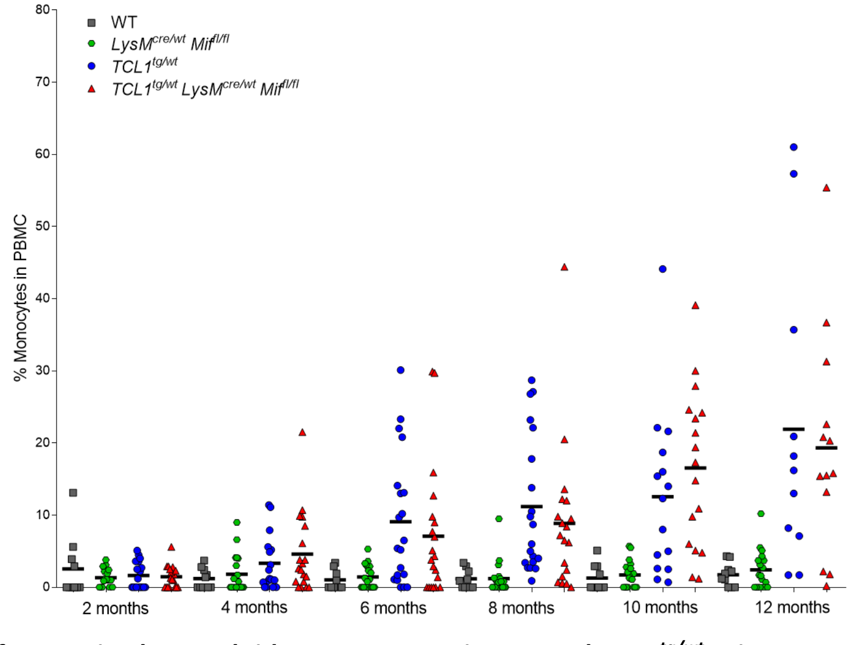


**Figure S1. No difference in the myeloid compartment in WT and $TCL1^{tg/wt}$ mice upon conditional Mif knockout in macrophages.**

A: Flow cytometric analysis of $CD11b^{+}$ myeloid cells in the peripheral blood of WT, $LysM^{cre/wt}$ $Mif^{fl/fl}$, $TCL1^{tg/wt}$ and $TCL1^{tg/wt}$ $LysM^{cre/wt}$ $Mif^{fl/fl}$ mice over one year.

B: Flow cytometric analysis of $CD14^{+}$ monocytes in the peripheral blood of WT, $LysM^{cre/wt}$ $Mif^{fl/fl}$, $TCL1^{tg/wt}$, and $TCL1^{tg/wt}$ $LysM^{cre/wt}$ $Mif^{fl/fl}$ mice over one year.
